# Fecal microbiota and their association with heat stress in *Bos taurus*

**DOI:** 10.1101/2021.10.06.463378

**Authors:** Bartosz Czech, Joanna Szyda, Kai Wang, Hanpeng Luo, Yachun Wang

## Abstract

Humans have been influencing climate changes by burning fossil fuels, farming livestock, and cutting down rainforests, which has led to global temperature rise. This problem of global warming affects animals by causing heat stress, which negatively affects their health, biological functions, and reproduction. On the molecular level, it has been proved that heat stress changes the expression level of genes and therefore causes changes in proteome and metabolome. The importance of a microbiome in many studies showed that it is considered as individuals’ “second genome”. Physiological changes caused by heat stress may impact the microbiome composition. In this study, we identified fecal microbiota associated with heat stress that was quantified by three metrics – rectal temperature, drooling, and respiratory scores and represented by their Estimated Breeding Values. For this purpose, the 16S rRNA sequencing technique was used. We analyzed the microbiota from 136 fecal samples of Chinese Holstein cows through a 16S rRNA gene sequencing approach. Sequence data were processed using a pipeline involving QIIME2 software together with SILVA database. Statistical modeling was performed using a negative binomial regression. The analysis revealed the total number of 24 genera and 12 phyla associated with heat stress metrics. *Rhizobium* and *Pseudobutyrivibrio* turned out to be the most significant genera, while *Acidobacteria* and *Gemmatimonadetes* were the most significant phyla. Phylogenetic analysis revealed that that three heat stress indicators quantify different metabolic ways of animals’ reaction to heat stress. Other studies already identified that those genera had significantly increased abundance in mice exposed to stressor-induced changes. Moreover, identified microbiota significantly associated with heat stress measures were mostly mesophilic, so their association seems to be due to heat stress-induced secondary, metabolic changes, and not directly by temperature. Moreover, high fold changes of many genera suggest that they may be used as biomarkers for monitoring the level of heat stress in cattle.

## 1. Introduction

Global warming and the resulting long-term increase in temperatures are the main cause of heat stress in mammals [1]. Moreover, selection towards high production yield in livestock and associated with its high metabolic load is an additional factor that makes livestock especially prone to overheating. Heat stress negatively affects health, reproduction, and other biological functions [2]. Specifically, in dairy cattle, heat stress impedes milk production, welfare, and growth [3]. Unfortunately, the phenomenon of heat stress is common in current ages, and we should understand how its long-term susceptibility affects organisms. On the genomic level, heat stress is manifested by transcriptional and post-transcriptional regulation of heat stress-associated genes [4]. It is known that *Bos indicus* has greater heat tolerance than *Bos taurus* [5], which indicates a genetic component of heat resistance. A few genes responsible for thermotolerance in dairy cattle – HSF1, MAPK8IP1, and CDKN1B have been recently identified [6]. However, the effect of heat stress on animal-associated microbiotas is not well known. In cattle genomics, bacteria are the main cause of mastitis – one of the most prevalent diseases of dairy cattle [7]. In general, the composition of gut microbiota depends on multiple factors – genetic [8], dietary [9] and environmental [10].

Heat stress belongs to environmental factors that may change the composition of the microbiota. Studies in cattle reported microbial species which abundance depends on heat stress conditions. Chen and colleagues [11] reported the effect of heat stress on physiological characteristics and circulation levels of immune activity and the microbiome. In an experimental study Zhao and colleagues [12] identified bacterial species in the rumen microbiome associated with heat stress. In their study it was found that heat stress has no effect on both alpha and beta diversity, however the effect on the richness of microbiota was identified, especially significant increase in the abundance of *Streptococcus, Enterobacteriaceae, Ruminobacter, Treponema* and *Bacteroidaceae*. Sales and colleagues [13] in his study also reported that heat stress influenced microbiota in beef cattle rumen. Particularly, they found genera *Flavonifractor, Treponema, Ruminococcus*, and *Carnobacterium* significantly associated with heat stress. However, assessing the composition of microbiota in farm animals’ environments is important to study its association with heat stress under breeding conditions. Moreover, the categorization between heat stress and normal conditions is a simplification. The level of an animal’s heat stress is a continuous variable and thus can be assessed using quantitative metrics. This however implies non-standard statistical modeling of the association between heat stress traits and microbiome as compared to the experimental-based, case-control setup. In our study, we focused on the identification of bacteria associated with heat stress measured by drooling score, rectal temperature, and respiratory score, expressed by the estimated breeding values.

## 2. Material and methods

The material comprises 136 fecal samples of 136 Chinese Holstein cows, which were collected in 2017, 2018, and 2019 directly in herds belonging to Beijing Shounong Livestock Development Co., Ltd. The cows from the same year had been fed with exactly the same total mixed ration diet for over 1 month and the cows from different years were fed with different total mixed ration diets with small change; however, all diets were based on corn silage and concentrate, and all the cows were fed ad libitum.

Fecal samples were collected directly from the cow’s rectum using a method a bit similar to rectal inspection. Around 7 AM, before the new feed is provided to the cow, is the time point we selected to take fecal samples, as cows are calmer after a good rest during the night, samples are easier to keep during the morning cooler period in summer, also we can be more sure that cows are in similar digestion stage without stimulation from feed for a relatively long time, and feces accumulated in the cow’s rectum. By wearing a disposable plastic long-armed glove, the sampler inserts his hand and arm into the cow’s rectum, first removed the outer part of feces accumulated in the rectum, and then grabbed a certain amount of feces from the inner part by hand, after a few feces mixing actions in the rectum. A disposable plastic long-armed glove can be used once for each cow. After the sampler’s hand holding feces moves out of the cow’s rectum, one can turn the glove outside in. Feces will naturally accumulate into the finger parts of the glove. By cutting a small hole at the tip of the finger parts of the glove, a fecal sample can be easily transferred into a properly labeled sterile 5 ml cryopreservation tube. Since big particles within feces may precipitate at the bottom of the finger parts of the glove, the very first part of the fecal sample can be discarded. Fecal samples are normally then placed on dry ice maximum for 3-4 h before they stored at -80°C at the laboratory.

The thermal environment during the sampling process was measured by temperature, humidity, and Temperature Humidity Index (THI) presented in Table 1.

**Table 1:** Characteristic of the thermal environment.

| sampling date | Temperature (Td) | Humidity (RH) | THI |
| --- | --- | --- | --- |
| 15 August, 2017 | 26.58 | 0.81 | 77.46 |
| 14 August, 2018 | 27.08 | 0.82 | 78.52 |
| 27 July, 2019 | 31.64 | 0.60 | 82.01 |

The procedures of DNA extraction, amplification, and sequencing were completed by Wekemo Tech Co., Ltd. (Shenzhen, China). Microbial DNA was extracted from fecal samples using the E.Z.N.A. soil DNA Kit (Omega Biotek, Norcross, GA, U.S.) according to manufacturer’s protocols. The final DNA concentration and purification were determined by NanoDrop 2000 UVvis spectrophotometer (Thermo Scientific, Wilmington, USA), and DNA quality was checked by 1% agarose gel electrophoresis. The V3-V4 hypervariable regions of the bacteria 16S rRNA gene were amplified with primers 338F(5’-ACTCCTACGGGAGGCAGCAG-3’) and 806R(5’-GGACTACHVGGGTWTCTAAT-3’) (for samples picked in 2017) as well as 341F(5’-CCTAYGGGRBGCASCAG-3’) and 806R(5’-GGACTACNNGGGTATCTAAT-3’) (for samples picked in 2018 and 2019) by thermocycler PCR system (GeneAmp 9700, ABI, USA). The PCR reactions were conducted using the following program: 3 min of denaturation at 95 °C, 27 cycles of 30 s at 95 °C, 30s for annealing at 55 °C, and 45s for elongation at 72 °C, and a final extension at 72 °C for 10 min. PCR reactions were performed in triplicate 20 *µL* mixture containing 4 *µL* of 5 × FastPfu Buffer, 2 *µL* of 2.5 mM dNTPs, 0.8 *µL* of each primer (5 *µM*), 0.4 *µL* of FastPfu Polymerase and 10 ng of template DNA. The resulted PCR products were extracted from a 2% agarose gel and further purified using the AxyPrep DNA Gel Extraction Kit (Axygen Biosciences, Union City, CA, USA) and quantified using QuantiFluor™-ST (Promega, USA) according to the manufacturer’s protocol. Amplicons were sequenced using the HiSeq-PE250 (samples picked in 2017) and MiSeq-PE300 (samples picked in 2018 and 2019) Illumina platforms in paired-end modes. Part of the sequence data analyzed previously by Zhang and colleagues [14] was used in this study.

All the heat stress phenotypes were measured as it was described by Luo and colleagues [15]. In order to correct for the environmental effects that may affect phenotype values, cows’ response to heat was expressed by breeding values for rectal temperature (RT), drooling (DS), and respiratory scores (RS) estimated using the following model:

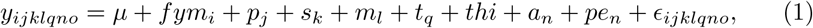

where *y*_*ijklqno*_ refers to phenotype (RS, DS or RT), *µ* is the population mean, *fym*_*i*_ is the fixed effect for farm-year, *p*_*j*_ is the fixed effect of parity, *s*_*k*_ is the fixed effect of lactation stage, *m*_*l*_ is the fixed effect of the indication if the animal is before or after milking, *t*_*q*_ is the fixed effect of testing time (morning or afternoon), *thi* is the fixed effect of temperature-humidity index, *a*_*n*_ is the animal additive genetic effect, *pe*_*n*_ is the permanent environmental effect, and *E*_*ijklqno*_ is the random residual. The covariance matrix of random effects has the following structure:

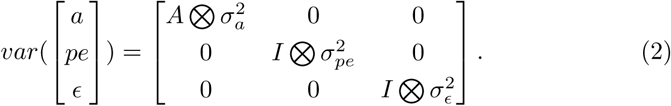

The total number of 155 cows were used to estimate the breeding values. The reliability of calculate EBVs was presented in Table 2.

**Table 2:**
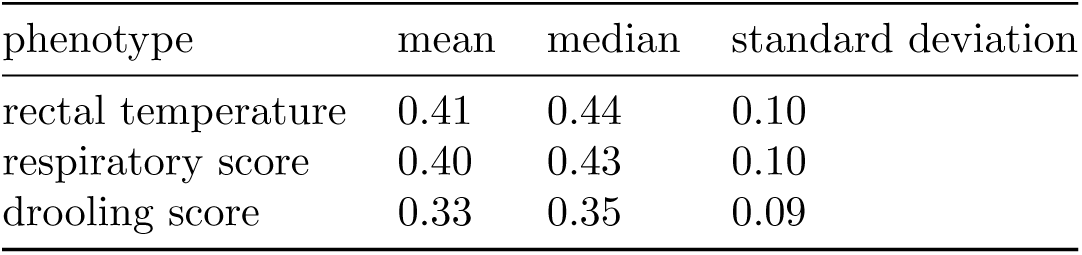
Descriptive statistics of the reliability of the estimated breeding values.

| phenotype | mean | median | standard deviation |
| --- | --- | --- | --- |
| rectal temperature | 0.41 | 0.44 | 0.10 |
| respiratory score | 0.40 | 0.43 | 0.10 |
| drooling score | 0.33 | 0.35 | 0.09 |

Furthermore, the breeding values were additionally corrected by deregression [16] in order to remove the ancestral information from the EBVs.

### 2.1. Ethics Committee Approval

The data collection process was carried out in strict accordance with the protocol approved by the Animal Welfare Committee of the China Agricultural University.

### 2.2. Processing of sequencing data

The first step of the analysis included quality control of sequenced data. For this purpose, the FastQC [17] software was used. Then, poor quality reads and adapter sequences were removed using Trim Galore [18]. Followingly, cleaned reads were processed using the QIIME 2 [19] software. First of all, data were dereplicated – reads that are 100% the same will be pooled together. Next, reads are were denoised – reads that occur very rarely are considered as PCR errors are removed as well as chimeric sequences and singletons. Those steps were done using DADA2 algorithm [20]. implemented in QIIME 2. Afterward, the Amplicon Sequence Variants (ASVs) table that represents counts of occurrence of a given sequence in a sample was created. Diversity within samples (*α*-diversity) was calculated using Simpson’s evenness and Shannon’s diversity indices using the phyloseq [21] R package. The association of microbes composition with heat stress factors was tested using aGLMM-MiRKAT test implemented in GLMM-MiRKAT R package [22].

The SILVA database [23] was used to classify ASVs taxonomically. For the classification, the naive Bayes algorithm implemented in scikit-learn Python package was used [24].

Since taxa originally assigned by the SILVA database represent different levels of taxonomy, they were aggregated to genera and phyla levels. Genera and phyla with a variance below one and that occurred in less than three samples were excluded from downstream analyses. Such filtered genera and phyla tables were used for the further differential abundance analysis. Additionally, for organoleptic testing of batch effect occurrent, the Uniform Manifold Approximation and Projection (UMAP; [25]) dimensional reduction technique was used to find potential sources of unwanted variability. The phylogenetic tree was generated using the align-to-tree-mafft-fasttree pipeline implemented in QIIME2 software.

### 2.3. Differential abundance analysis

The edgeR [26] R package was used for the normalization of the processed ASV table as well as for statistical modeling of the association between the abundance of microbiota and heat stress indicators. In particular, the Trimmed Mean of M-values (TMM) based normalization [27] was applied. It identified and excluded highly abundant and highly variables genera and phyla, whereupon weighted mean of an abundance of remaining groups were used for the actual normalization [28]. The association between genera/phyla abundance and heat stress indicators was modeled using the negative binomial distribution:

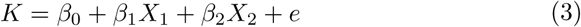

where *K* represents the counts of reads for a given genus/phylum, *β*_0_ is the intercept, *β*_1_ is the effect of the DRP (expressed by log fold change), *X*_1_ is the design matrix for DRP, *β*_2_ is the effect of the sampling year class, *X*_2_ is the incidence matrix for sampling year, *e* is the random error.

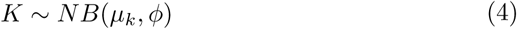

where *µ*_*k*_ represents the mean of counts reads, and *φ* is the dispersion parameter such that 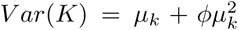 calculated using Cox-Reid approximate conditional inference moderated towards the mean [29].

The significance of the effect of DRP on the relative abundance of the genus/phylum was tested using the Likelihood Ratio Test [30]. The test statistic is as follows:

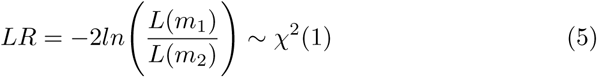

where *m*_1_ is the reduced model (i.e. the formula (5) without the effect of DRP), and *m*_2_ is the full model (5).

Since each genus/phylum is tested separately, multiple testing correction method was applied using a False Discovery Rate (FDR) [31].

## 3. Results

### 3.1. Processing and classification of sequence variants

46,825 unique sequences of V3 and V4 regions with a total of 6,486,706 reads were identified and classified. Table 3 summarizes the classification of ASVs based on the different taxonomic levels. In general, reads were classified into two domains – archaea (0.01%) and bacteria (99,99%). Almost all reads could be taxonomically assigned up to order, but species could be assigned only to 2.35% of reads. Further analysis was carried out using genus-level and phylum-level resolution.

**Table 3:** Amplicon Sequence Variants classification results.

| Taxonomic level | Number of unique features | Percent of classified reads |
| --- | --- | --- |
| Domain | 2 | 100.00 |
| Phylum | 29 | 97.94 |
| Class | 72 | 97.81 |
| Order | 114 | 97.50 |
| Family | 156 | 70.16 |
| Genus | 235 | 20.93 |
| Species | 152 | 2.35 |

#### 3.1.1. Microbiota composition

The general composition of microbiota presented in all samples was presented on bar plot using genus-level and phylum-level resolution. Figure 1 presents the relative abundance of genera with average proportions of more than 0.5%. We can see that *Clostridium* is a genus with the highest relative abundance (15.14%). There were 209 genera with a relative abundance of less than 0.5%. Figure 2 presents the analogous visualization for phyla. *Firmicutes* is a phylum with the highest relative abundance (63.66%). Regarding the less abundant phyla, there were identified 22 phyla with less than 0.5% of the relative abundance.

**Figure 1:**
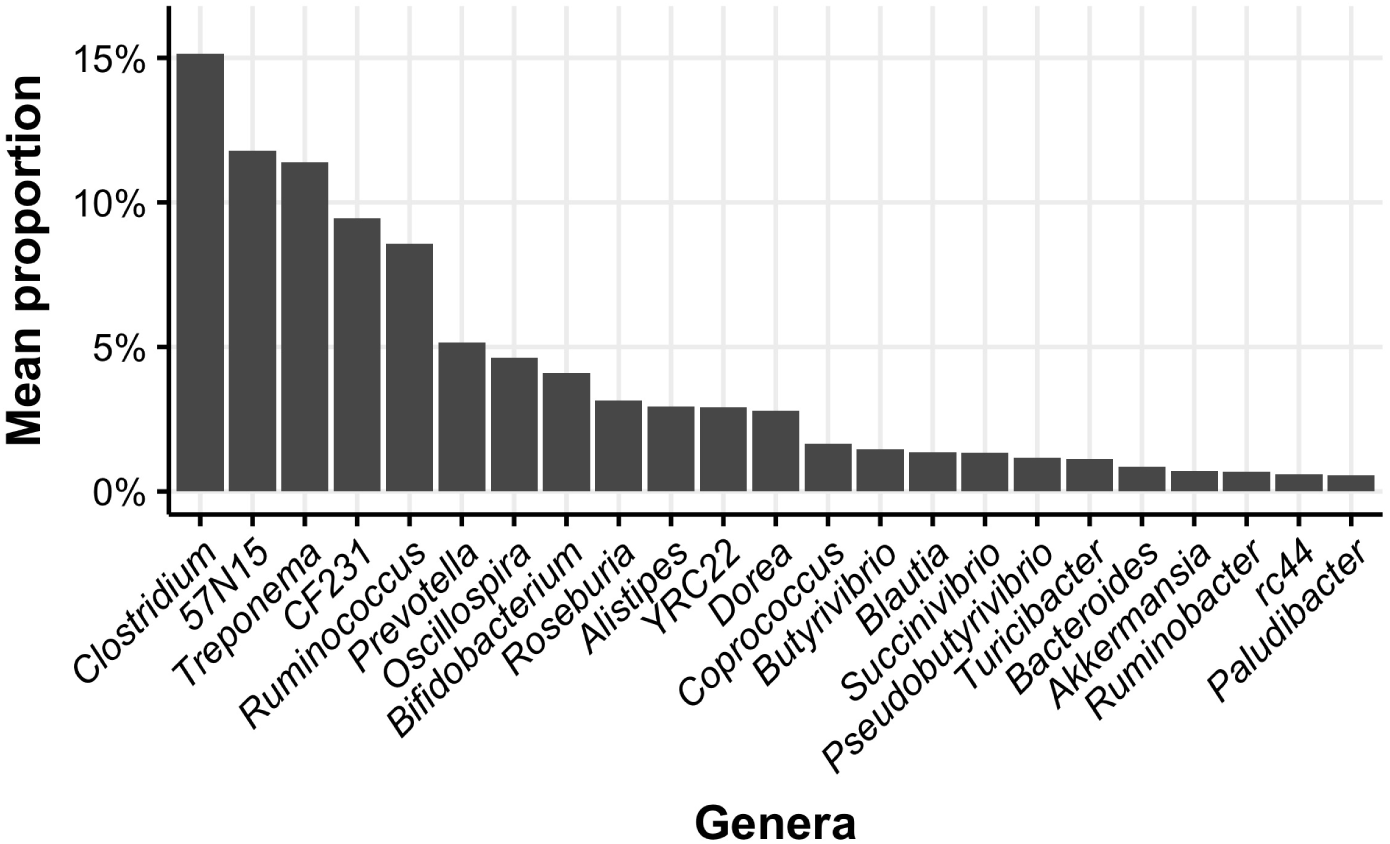
The relative abundance of genera with average proportions of more than 0.5%.

**Figure 2:**
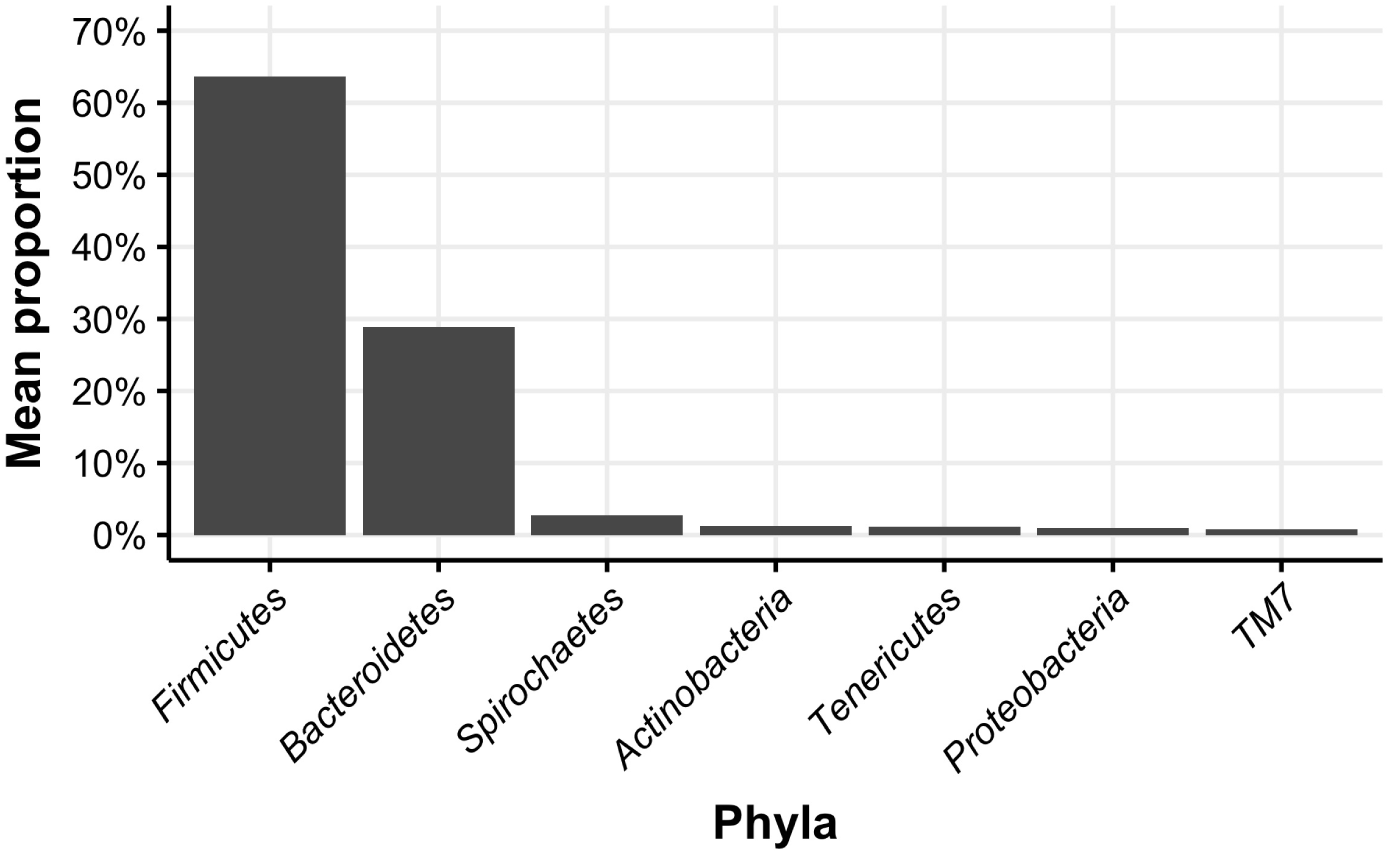
The relative abundance of phyla with average proportions of more than 0.5%.

#### 3.1.2. Clustering

Genera table was then clustered using UMAP algorithm. The projection of the UMAP coordinates calculated from the ASVs counts matrix on the genus level demonstrates three distinct clusters (Figure 3), which reflect the sampling year. In further analysis, the effect of the sampling year was corrected.

**Figure 3:**
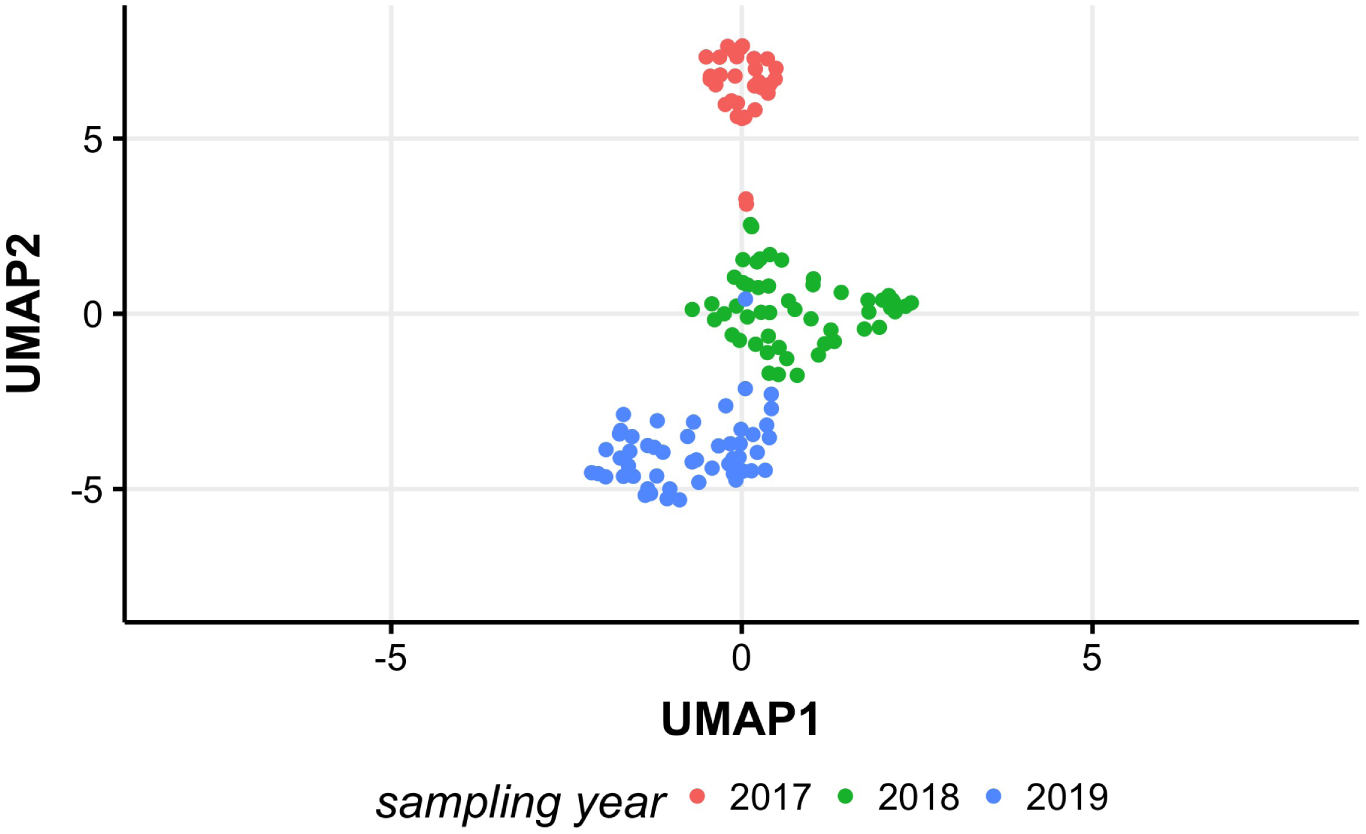
UMAP projection of the ASVs counts matrix on the genus level.

### 3.2. Correlation analysis of diversity metrics

In order to check whether the general diversity of microbiota within samples is correlated with the DRPs, a correlation analysis was performed. Correlations were generally positive, but non-significant (Table 4) with the highest correlation estimated between DRPs for the respiratory score and the Simpson’s evenness index (0.27). Overall, non-significant correlations indicate that there is no linear dependence between DRPs and sample diversity calculated based on the abundance of genera.

**Table 4:** Pearson correlation coefficients between DRPs and alpha diversity measures expressed by Simpson’s evenness and Shannon diversity.

| DRP | Simpson's evenness | Shannon diversity |
| --- | --- | --- |
| Rectal temperature | 0.25 | -0.04 |
| Drooling score | 0.13 | 0.23 |
| Respiratory score | 0.27 | 0.11 |

### 3.3. Relationship of EBVs with microbiomes composition

aGLMM-MiRKAT test was calculated to test the association between the microbial community composition and EBVs. None of the analyzed EBVs showed statistically significant association with the microbial composition. It means that individual genera and phyla should be considered in a statistical model.

### 3.4. Differential abundance analysis

Based on the results of the negative binomial model and considering *FDR* ≤ 0.05 22 genera were significantly associated with rectal temperature with all but one (*Helococcus*) of them showing decreased abundance with the increase of rectal temperature. *Rhizobium* – that represents soil bacteria – was the most associated genus with the rectal temperature. The occurrence of this bacteria might be observed perhaps due to the specific metabolism or the specific plant diet.

Succinivibrio was the only genus associated with respiratory score and *Pseudobutyrivibrio* – with the drooling score. There was no overlap between genera significant for the three heat stress indicators (Table 5). Differential abundance analysis of phylum showed that 6 phyla were significiantly associated with rectal tempearture. All of them showed decreased abundance with the increase of rectal temperature. *Fibrobacteres* was the only phylum associated with respiratory score. Surprisingly, for drooling score, 5 differentially abundant phyla were identified. Five of them showed increase abundance with the increase of rectal temperature. Only *Fibrobacteres* showed the descrease abundance. *Fibrobacteres* was significantly associated with both drooling and respiratory scores, while *Nitrospiarae, Gemmatimonadetes, Acidobacteria*, and *Planctomycetes* were significantly associated with both rectal temperature and drooling score.

**Table 5:**
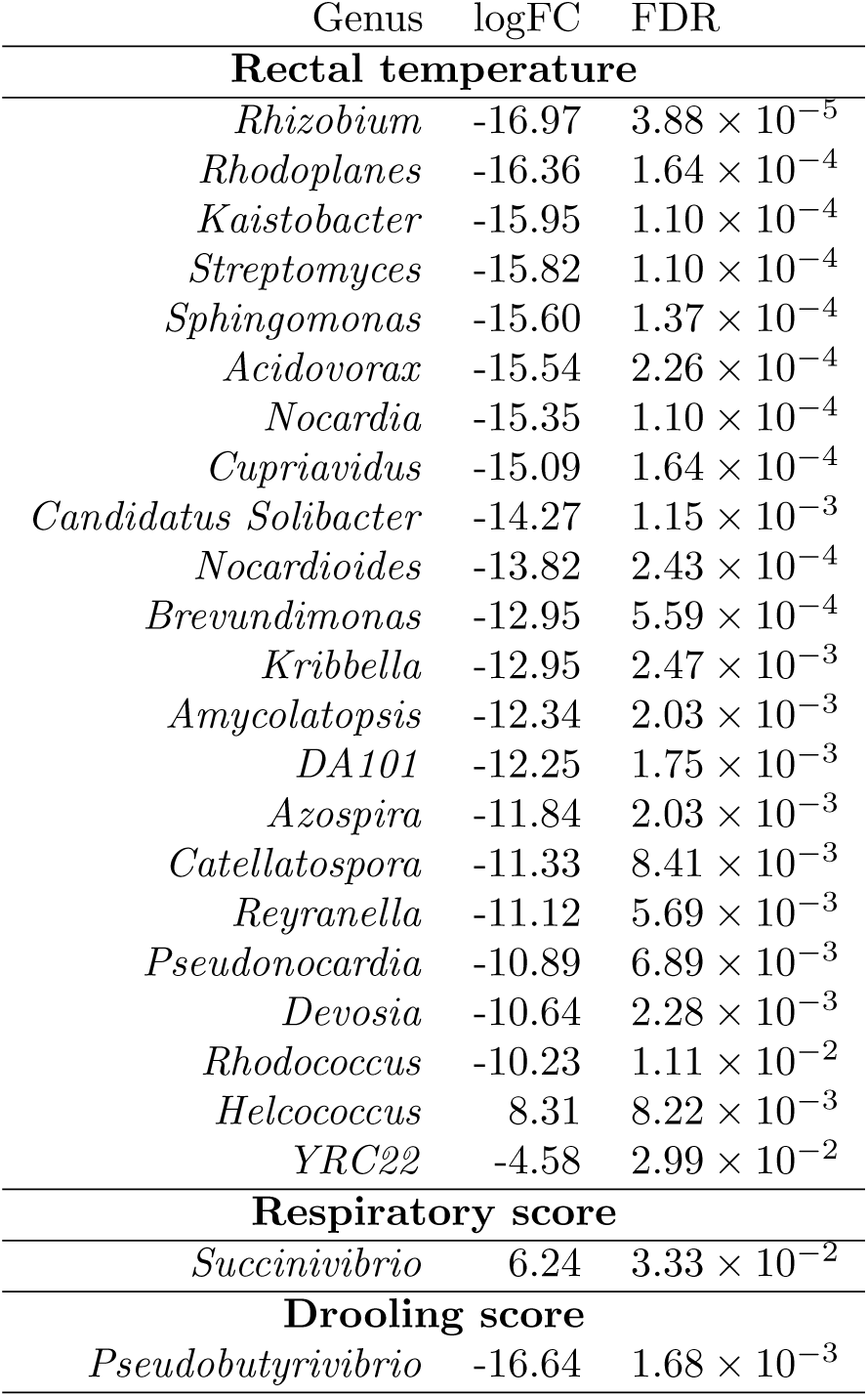
Significant differentially abundant genera.

In order to check the genetic relationship between those associated genera, the phylogenetic tree was created based on the 16S rRNA sequences. The genetic relationship of the associated genera was shown in Figure 4. Colors indicate the association between the genera and the phenotype. We can see, that *Succinivibrio* that was associated with the respiratory score phenotypes created the single clade with all the genera associated with the rectal temperature. Only *Pseudobutyrivibrio* that was associated with the drooling score creates a single, separate clade.

**Figure 4:**
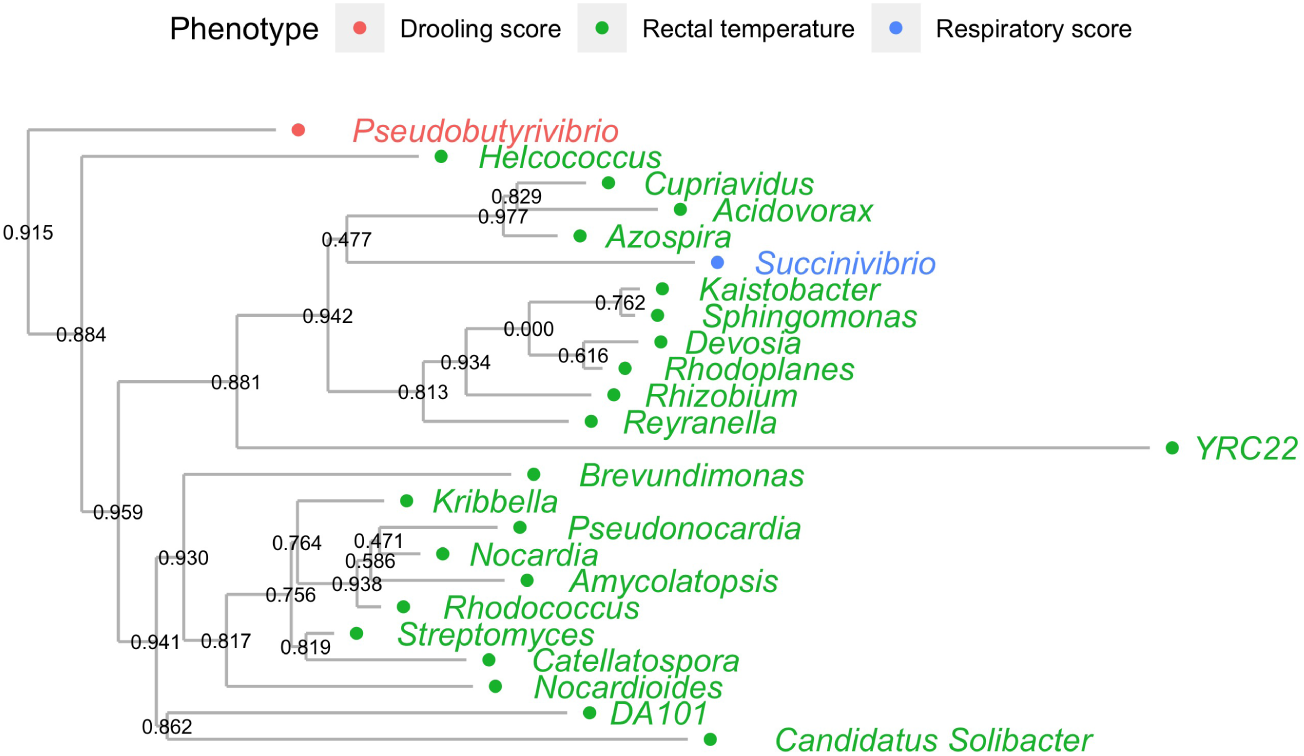
Phylogenetic tree of genera identified in fecal samples. Color indicates significantly associated genera with a given phenotype.

## 4. Discussion

This study aimed to identify genera that are associated with the rectal temperature, drooling score, and respiratory score, and in the consequences, associated with heat stress. The quantitative pseudophenotypes were used in order to model animals’ microbiomes under conventional production conditions, without setting up a case (heat stress conditions) – control (standard conditions) experiment. Such an approach allows for the estimation of genera effect on heat stress under real conditions underlying dairy herd management.

The general composition of microbiota was not altered by heat stress. Therefore we focussed on single genera as potentially involved in heat stress response. Most of the genera were significantly associated with rectal temperature which might be caused by the fact that samples and measurement came from the same environment (rectum). Since most of the significantly associated genera showed decreased abundance with the increase of heat stress, we can assume, that heat stress favors the inhibition of growth of some microbial populations.

**Table 6:** Significant differentially abundant phyla.

| Phylum | logFC | FDR |
| --- | --- | --- |
| **Rectal temperature** |  |  |
| *Acidobacteria* | -26.99 | $1.18 \times 10^{-16}$ |
| *Gemmatimonadetes* | -22.40 | $5.03 \times 10^{-13}$ |
| *Chloroflexi* | -21.07 | $2.98 \times 10^{-11}$ |
| *Nitrospirae* | -15.19 | $1.18 \times 10^{-7}$ |
| *Planctomycetes* | -11.19 | $5.52 \times 10^{-5}$ |
| *Euryarchaeota* | -6.80 | $2.99 \times 10^{-2}$ |
| **Respiratory score** |  |  |
| *Fibrobacteres* | -8.67 | $2.58 \times 10^{-2}$ |
| **Drooling score** |  |  |
| *Fibrobacteres* | -16.72 | $5.22 \times 10^{-4}$ |
| *Nitrospirae* | 15.73 | $4.22 \times 10^{-4}$ |
| *Gemmatimonadetes* | 15.17 | $4.69 \times 10^{-4}$ |
| *Acidobacteria* | 14.46 | $4.71 \times 10^{-4}$ |
| *Planctomycetes* | 12.50 | $1.13 \times 10^{-2}$ |

Based on the current literature, Bailey [32] observed a reduced abundance of bacteria in genus *Pseudobutyrivibrio* in mice exposed to stressor-induced changes. Such reduced abundance was also observed by us the association with a drooling score. Baek [33] in his study observed that *Succinivibrio* shows increased abundance in cows under heat stress. In our study, this genus was also associated with the respiratory score metric. Interestingly, *Helcococcus*, the only genus that abundance increased with increasing rectal temperature, has not been reported in studies focused on heat stress and any stress-induced conditions, but it was reported as associated with postpartum endometritis by Miranda CasoLuengo [34]. Moreover, [35] showed that *Streptomyces* was reported as a genus with enriched relative abundance in Jersey cows in the normal condition compared to the heat stress condition. It is worthwhile to mention that many genera reported with the association to the rectal temperatures show the high fold change, suggesting that increased rectal temperature has a high impact on microbiota composition. *Proteobacteria* phylum that represents most of the associated genera in our study seems to be the most important phylum in heat stress conditions. Yu [36] already reported that *Proteobacteria* and *Firmicutes* are the most common phyla associated with heat stress conditions. Interestingly, analysis based on the phylum resolution showed that there were overlapping phyla. *Fibrobacteres* turned out to be the significantly associated phylum with respiratory and drooling scores. This phylum was already reported as significant in heat stress analysis of pigs reported by He [37]. *Chloroflexi* and *Planctomycetes* significantly associated with rectal temperature were also reported as a significant phyla in the analysis of short-term acute heat stress on the rumen microbiome of Hanwoo steers [33].

Differences found in microbial compositions and in genera/phyla abundance suggest that those changes might occur due to adapting to climate change. In this study, the abundance of *Fibrobacteres* was decreased due to heat stress. The role of this bacteria is the degradation of plant-based cellulose in ruminants and acetate production. Ransom-Jones and colleagues [38] reported that glycosyl hydrolases of *Fibrobacteres* may produce carbohydrate activators, including cellulose enzymes and in consequence, cows may produce more energy with acetate in the rumen that can be associated with heat production. Some bacteria (e.g. *Pseudobutyrivibrio*) were described as a part of the microbiome, but their impact on host physiology is not yet known.

Heat stress modeled as a binary variable (i.e. normal vs. stress conditions) provides valuable insights into the understanding of the microbiome association to heat stress, however, it should be beard in mind that the real, production environment of a dairy cow markedly deviates from the experimental conditions. The most obvious differences comprise duration, intensity, and variation in ambient temperatures, which are typically not modeled in experiments. Therefore, our study, despite being more challenging from the analytical perspective, provided an attempt to analyze the microbiome dynamics directly in a production herd. In such a situation, an important aspect of the analysis is the heat stress “phenotype”. In order to pre-correct for a whole series of genetic (i.e. familial relationship) and environmental effects (such as parity or lactation stage) possibly affecting the heat stress indicator measurements, prior to the actual heat stress modeling, we decided to use breeding values as pseudophenotypes, which were then deregressed in order to remove ancestral and familiar contributions. Such an approach provided a novel approach for the investigation of bacteria in dairy cattle under heat stress condition.

## Author Contributions

B.C., Y.W. and J.S conceived and conducted the experiment, B.C. analyzed the results and wrote the manuscript in consultation with J.S., K.W., H.L. and Y.W. All authors reviewed the manuscript.

## Acknowledgments

Calculations have been carried out using resources provided by Wroclaw Centre for Networking and Supercomputing (http://wcss.pl), grant No. 509. Computations were carried out at the Poznan Supercomputing and Networking Centre

## Data Availability Statement

The sequence data are available in the NCBI Sequence Read Archive with accession number SRP202074. Other datasets generated and/or analyzed during the current study are not publicly available due to institutional constraints but are available from Yachun Wang on reasonable request.

